# Prelude to Malignancy: A Gene Expression Signature in Normal Mammary Gland from Breast Cancer Patients Suggests Pre-tumorous Alterations and Is Associated with Adverse Outcomes

**DOI:** 10.1101/2024.02.07.579074

**Authors:** Maria Andreou, Marcin Jąkalski, Katarzyna Duzowska, Natalia Filipowicz, Anna Kostecka, Hanna Davies, Monika Horbacz, Urszula Ławrynowicz, Katarzyna Chojnowska, Bożena Bruhn-Olszewska, Jerzy Jankau, Ewa Śrutek, Manuela Las-Jankowska, Dariusz Bała, Jacek Hoffman, Johan Hartman, Rafał Pęksa, Jarosław Skokowski, Michał Jankowski, Łukasz Szylberg, Wojciech Zegarski, Magdalena Nowikiewicz, Tomasz Nowikiewicz, Jan P. Dumanski, Jakub Mieczkowski, Arkadiusz Piotrowski

## Abstract

Despite advances in early detection and treatment strategies, breast cancer recurrence and mortality remain a significant health issue. Recent insights suggest the prognostic potential of microscopically healthy mammary gland, in the vicinity of the breast lesion. Nonetheless, a comprehensive understanding of the gene expression profiles in these tissues and their relationship to patient outcomes is still missing. Furthermore, the increasing trend towards breast-conserving surgery may inadvertently lead to the retention of existing cancer-predisposing mutations within the normal mammary gland. This study assessed the transcriptomic profiles of 242 samples from 83 breast cancer patients with unfavorable outcomes, including paired uninvolved mammary gland samples collected at varying distances from primary lesions. As a reference, control samples from 53 mammoplasty individuals without cancer history were studied. A custom panel of 634 genes linked to breast cancer progression and metastasis was employed for expression profiling, followed by whole-transcriptome verification experiments and statistical analyses to discern molecular signatures and their clinical relevance. A distinct gene expression signature was identified in uninvolved mammary gland samples, featuring key cellular components encoding keratins, CDH1, CDH3, EPCAM cell adhesion proteins, matrix metallopeptidases, oncogenes, tumor suppressors, along with crucial genes *(FOXA1, RAB25, NRG1, SPDEF, TRIM29*, and *GABRP*) having dual roles in cancer. Enrichment analyses revealed disruptions in epithelial integrity, cell adhesion, and estrogen signaling. This signature, named KAOS for Keratin-Adhesion-Oncogenes-Suppressors, was significantly associated with reduced tumor size but increased mortality rates. Integrating molecular assessment of non-malignant mammary tissue into disease management could enhance survival prediction and facilitate personalized patient care.

## INTRODUCTION

Breast cancer remains a pervasive global health concern and represents the most prevalent malignancy worldwide, surpassing lung cancer with 2.26 million reported incidents in 2020 ^1,2^. Improved mammographic screening and widespread educational initiatives, resulting in increased self-monitoring, have facilitated the early detection of breast carcinomas in asymptomatic stages. Consequently, breast-conserving surgery (BCS) has become increasingly prominent as a favored treatment approach, involving the removal of the tumor, while minimizing the removal of healthy tissue, thereby preserving a substantial portion of the breast ^3,4^. However, despite histopathologically negative surgical margins, suggesting a complete tumor excision during BCS, a considerable proportion of patients experience recurrence rates as high as 19.3% in those receiving radiotherapy and 35% in patients subjected to BCS alone ^5^, while administration of adjuvant chemotherapy reportedly reduces the recurrence rate by one-third ten years post-surgery ^6^. The underlying cause of recurrence, whether it is due to undetected residual disease or the development of additional changes within the unexcised mammary gland remains unclear.

Currently, decisions regarding therapeutic management primarily rely on pathological examination and genetic tests performed solely on fragments originating from tumor as well as resection margins (mammary gland tissue in the tumor perimeter excised during surgery). However, emerging evidence suggests that the normal mammary gland tissue surrounding the cancerous lesion holds promise of prognostic value ^7–10^. Notably, the inclusion of normal, cancer-adjacent tissue samples in study designs, significantly enhances the accuracy of overall survival predictions compared to relying solely on tumor data ^11^. While previous studies have investigated paired normal and cancerous tissue samples ^12–15^, the association of transcriptomic landscape of normal, uninvolved mammary gland tissue, located at greater distance from the primary tumor and remaining in the patient’s body following BCS, with unfavorable patient outcomes has not been thoroughly investigated. Furthermore, the limited availability of an adequate number of control samples in study designs posed challenges in interpreting findings. Taking these factors into account, our study aimed to investigate a unique cohort of breast cancer patients characterized by adverse prognosis, with comprehensive follow-up data that extended to nearly a decade after their initial surgeries. We employed targeted RNA sequencing, to analyze the transcriptomic profiles of primary tumor and paired proximal and distal uninvolved mammary gland samples, as well as mammary glandular tissue samples from control individuals without any personal and familial history of cancer.

Our custom RNA-seq panel effectively discriminates between malignant (primary tumor) and non-malignant (uninvolved margin and control) tissues, by elucidating unique gene expression patterns for each group. Notably, our study uncovers the existence of a pre-tumorigenic microenvironment within the seemingly normal mammary gland tissue, displaying an association with smaller tumor size and higher patient mortality.

## MATERIALS AND METHODS

### Patient recruitment, sample collection and RNA isolation

We analyzed specimens obtained from 83 individuals who had been diagnosed with breast cancer including 70 who underwent breast-conserving surgery and 13 with mastectomies. All recruited individuals did not receive neoadjuvant therapy and were characterized by the presence of recurrent disease (metastasis to the breast or secondary organs) and/or the appearance of a second independent tumor and/or death in the following 10 years (Table 1, Additional File 1, Supplementary Table 1). For 2 individuals, two distinct samples from multifocal primary tumor (described as PT1 and PT2) were obtained. Fifty-three individuals subjected to breast reduction surgery without personal and familial history of cancer were recruited as controls (Table 1, Additional File 1, Supplementary Table 2).

**Table 1.** Summarized clinicopathological characteristics of breast cancer patient and control cohort. PT, UMD, and UMP samples were collected from 83 individuals diagnosed with breast cancer. CTRL samples were collected from 53 individuals without any personal and familial history of cancer. Histological evaluation was performed to identify tumor samples and confirm the normal histology of uninvolved margin and control samples. PT samples were classified as Invasive Ductal Carcinoma (IDC), Invasive Lobular Carcinoma (ILC), mixed (ICD-ILC), or other. Estrogen (ER), progesterone (PR), and ERBB2 (HER2) receptors were evaluated based on immunostaining. Biological subtypes were assigned based on ER/PR/HER2 and Ki67 status.. Recurrent disease was reported for 33 patients, the presence of a second, independent tumor was confirmed for 31 patients, and 50 patients died by the time of last contact. Detailed clinicopathological information is provided in Additional File 1, Supplementary Table 1.

|  |  |
| --- | --- |
| **Number of individuals** | 136 |
| Breast cancer (BC) patients | 83 |
| Controls (CTRL) | 53 |
| **Age (median/range)** |  |
| BC patients | 62 (23-85) |
| CTRL | 44 (18-76) |
|  | p.value= 1.324e-09 |
| **Collected samples (BC patients)** | 242 |
| Primary tumor, PT | 79 |
| Uninvolved margin distal from PT, UMD | 81 |
| Uninvolved margin proximal to PT, UMP | 82 |
| **Histology (BC patients)\*** |  |
| Invasive ductal carcinoma, IDC | 63 |
| Invasive lobular carcinoma, ILC | 4 |
| IDC - ILC | 7 |
| other | 9 |
| **Receptors (BC patients)\*** |  |
| Estrogen, ER (positive/negative) | 62 / 21 |
| Progesterone, PR (positive/negative) | 48 / 35 |
| HER2 (positive/negative) | 17 / 61 |
| **Subtype\*** |  |
| Luminal A | 16 |
| Luminal B | 40 |
| HER-2 enriched | 9 |
| Triple-negative | 12 |
| Not available | 6 |
| **Follow-up information (BC patients)** |  |
| Recurrence (yes/no) | 50 / 33 |
| Second cancer (yes/no) | 31 / 52 |
| Death (yes/no) | 50 / 33 |

Written informed consent was obtained from all enrolled individuals. The study was approved by the Bioethical Committee at the Collegium Medicum, Nicolaus Copernicus University in Toruń (approval number KB509/2010) and by the Independent Bioethics Committee for Research at the Medical University of Gdansk (approval number NKBBN/564/2018 with amendments). Graphical representation of the project workflow can be found in Figure 1A. A total of 295 samples, including primary tumor (PT), uninvolved mammary gland distal from (UMD, 1.5-5 cm), and proximal to the primary tumor (UMP, at least 1 cm from the primary tumor and always in shorter distance than UMD), as well as normal mammary gland from control individuals (CTRL), were collected in the Oncology Centre in Bydgoszcz, the University Clinical Centre in Gdańsk, and Karolinska Institute and deposited, along with clinical data and follow-up information, in biobank of our unit at the Medical University of Gdansk ^16^. PT, UM, and CTRL samples collected were frozen at -80°C. Detailed sampling design is presented in Figure 1B. All fragments prepared for molecular analysis were microscopically evaluated to identify tumor fragments and confirm normal histology of uninvolved margins and controls. RNA was extracted from tissues using the RNeasy Mini according to the original protocol with two modifications: (i) 1-bromo-3-chloropropane was used instead of chloroform to prevent foaming and emulsification and (ii) the elution was carried out with 90 µl of water for PT and 30 μl of water for UM and CTRL samples, followed by repeated elution with the entire volume of the original eluate (Qiagen, Germantown, MD). RNA concentration and quality were determined using Agilent TapeStation (Agilent Technologies).

**Figure 1.**
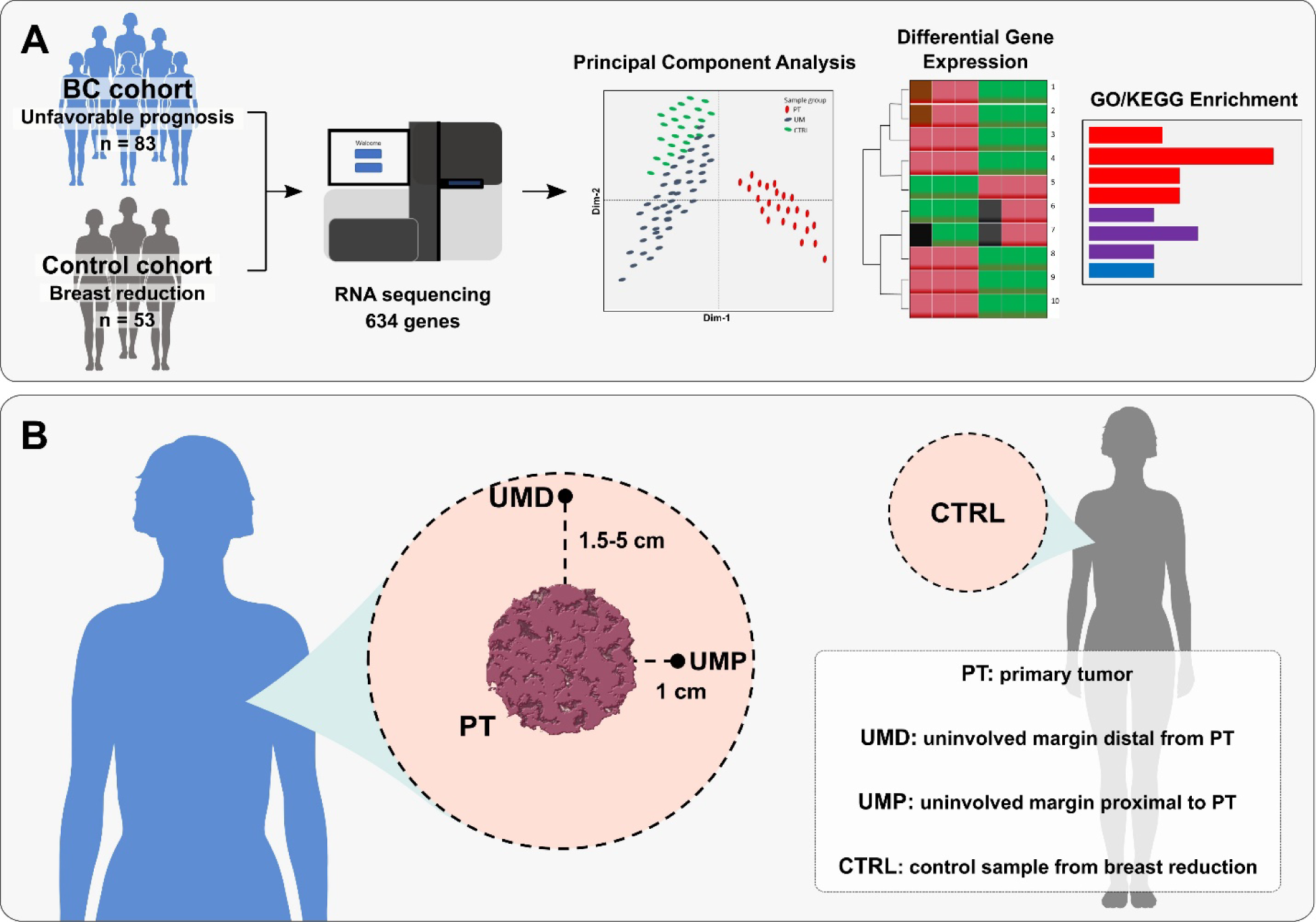
**A) Graphical representation of the project workflow**. A total of 295 fresh-frozen primary tumor (PT), uninvolved mammary gland excised proximal (UMP, >1 cm) and distal (UMP, 1.5-5 cm) from the PT, and control samples (CTRL) were collected from 83 individuals diagnosed with breast cancer and 53 individuals subjected to breast reduction surgeries respectively. After RNA extraction and library construction, targeted RNA sequencing was performed using a customized panel including genes previously associated with breast cancer. Bioinformatics analysis implementing standard tools was used to investigate expression patterns in PT, UM, and CTRL samples, as well as associations with follow-up clinical information. **B) Detailed sampling design.** Tissue blocks were collected from PT, UMP, and UMD samples from breast cancer patients with unfavorable outcomes. UMP was always collected in smaller distance than UMD from the PT. Tissue samples were evaluated by two independent pathologists to confirm the normal histology of uninvolved margin (UM) and control (CTRL) samples and identify tumor areas in breast cancer cases. Control (CTRL) mammary gland samples were collected from individuals subjected to breast reduction surgeries without personal and familial history of cancer. Parts of the figure were drawn by using pictures from Servier Medical Art. Servier Medical Art by Servier is licensed under a Creative Commons Attribution 3.0 Unported License (https://creativecommons.org/licenses/by/3.0/).

### Targeted RNA sequencing

The targeted RNA sequencing panel, designed with Roche NimbleDesign online tool (Roche, now HyperDesign, https://hyperdesign.com/#/), covered 7,229 regions with a total length of 1,243,523 bp. The panel includes 634 genes selected from literature research (Additional File 1, Supplementary Table 3). The genes have been associated with breast cancer and processes related to its dissemination and metastasis, such as epithelial-to-mesenchymal transition, cell death, and apoptosis. Furthermore, the panel incorporated genes from the AIMS and PAM50 predictors, originally developed to classify breast tumors into five distinct subtypes: Luminal A, Luminal B, HER2 enriched, basal-like, and normal-like ^17,18^. Sequencing libraries were prepared using KAPA RNA HyperPrep kit (Illumina Platforms, KR1350-v2.16) and the BRAVO NGS workstation (Agilent) with the dedicated automatization protocol (KAPA RNA Hyperprep kit KR1350-v.1.16). Hybridization was carried out with SeqCap RNA Choice Probes using the KAPA HyperCapture Reagent and Bead kit (v2, Roche Sequencing Solutions, Inc.) according to SeqCap RNA Enrichment System User’s Guide (v.1.0) with slight modifications. Component A was replaced with formaldehyde, and the Multiplex Hybridization Enhancing Oligo Pool was replaced with Universal Blocking Oligos (UBO). Next, the cDNA libraries were quantified using the KAPA Library Quantification kit (KR0405-v11.20, Kapa Biosystems, Woburn, USA). Paired-end reads of 150 bp were generated using TruSeq RNA Access sequencing chemistry on HiSeq X instrument (Illumina, San Diego, CA) by an external service provider (Macrogen Europe, Amsterdam, The Netherlands). The sequencing coverage and quality statistics for each sample are summarized in Additional File 1, Supplementary Table 4.

### Data analysis

#### Processing of sequencing data from the targeted panel

The raw RNA-seq data were first subjected to quality check using FastQC (https://www.bioinformatics.babraham.ac.uk/projects/fastqc/, version 0.11.9), followed by adapter trimming with BBDuk from the BBTools package (https://sourceforge.net/projects/bbmap/, version 38.36) with the following set of parameters: ktrim=r, k=23, mink=11, hdist=1, minlen=70, tpe, tbo. The processed reads were then mapped against the reference human genome (hg38, GENCODE version 35) using STAR (version 2.7.3a) ^19^. The generated ReadsPerGene.out.tab files (obtained through –quantMode GeneCounts parameter) were used to extract raw read counts that mapped to the annotated genes. Subsequently, the merged raw read matrix was generated with a custom R script and further processed with the edgeR (version 3.36.0) ^20^. First, the genes were filtered to keep only those whose expression was at least 1 count per million (CPM) in at least one sample. Next, the processed gene expression matrix was normalized using the TMM method in edgeR ^21^.

Principal Component Analyses (PCA) were performed to identify outlier samples using the R package FactoMineR (version 2.4) ^22^. This was carried out in several rounds: for all sample types separately (controls, tumor samples, etc.) and for all samples merged.

### Cancer subtype prediction

PAM50 and AIMS classifiers were applied to normalized gene expression matrices using the R package genefu (version 2.26) ^17,18,23^. The samples were classified into one of the following categories: Normal, LumA, LumB, Her2, or Basal.

### Sample clustering and differential expression analyses

Gene expression heatmaps were generated with the pheatmap R library (version 1.0.12). Sample clusters were identified using the built-in hierarchical clustering function of pheatmap (default parameters), which uses the Euclidean distance as the similarity measure and the complete linkage method. The number of sample clusters was set to four.

EdgeR was used to identify differentially expressed genes (DEGs) using the Quasi-Likelihood F-test (QLF) with a significance threshold set to 0.05 (False Discovery Rate, FDR) (19). Differential expression (DE) analyses were performed in two modes: first, to perform pairwise sample type comparisons; second, to additionally include the information on cluster membership of the samples. We included age as a covariate in the DE analyses due to the significant age difference between controls (44, 18-76) and cancer patients (62, 23-85) (Wilcoxon test). The identified sets of differentially expressed genes were subjected to enrichment analyses of Gene Ontology (GO) terms and KEGG pathways using ClusterProfiler ^24^.

### External RNA-seq datasets

External bulk RNA-seq datasets were used to corroborate our findings. These datasets originated from other breast cancer study in our lab. The sample collection and processing methods were consistent with those described in this study. Their FASTQ files were processed from scratch using the same tools as described above. Data integration was done with the ComBat_seq function from the sva R library (version 3.42.0, default parameters) to adjust for the effect of different data batches ^25,26^.

### Code availability

Unless otherwise stated, all analyses were conducted in R (version 4.1.2). The code used for data processing is available on GitHub: https://github.com/jakalssj3/Breast_cancer_KAOS

## RESULTS

### Clear delineation between malignant and non-malignant tissues

Principal component analysis (PCA) of all samples, using the normalized expression profiles of panel genes, revealed distinct differences between malignant (PT) and non-malignant (UM + CTRL) samples, identified by the first principal component (Figure 2A). Fourteen outliers were identified and excluded from downstream analyses, leaving 295 samples (53 controls, 163 margins, and 79 tumor samples). Differential expression analysis, which used non-malignant samples as a baseline, showed the largest set of deregulated genes (FDR ≤ 0.05, log-fold change of ≥1) when comparing PT against all non-malignant tissues (CTRLs, UMs) (Figure 2B). The number of differentially expressed genes decreased when comparing PT tissues with CTRLs or UMs separately. Interestingly, a relatively small number of differentially expressed genes was found between UMP and UMD, suggesting similar expression profiles and minor effects regardless of their physical distance from the primary tumor.

**Figure 2.**
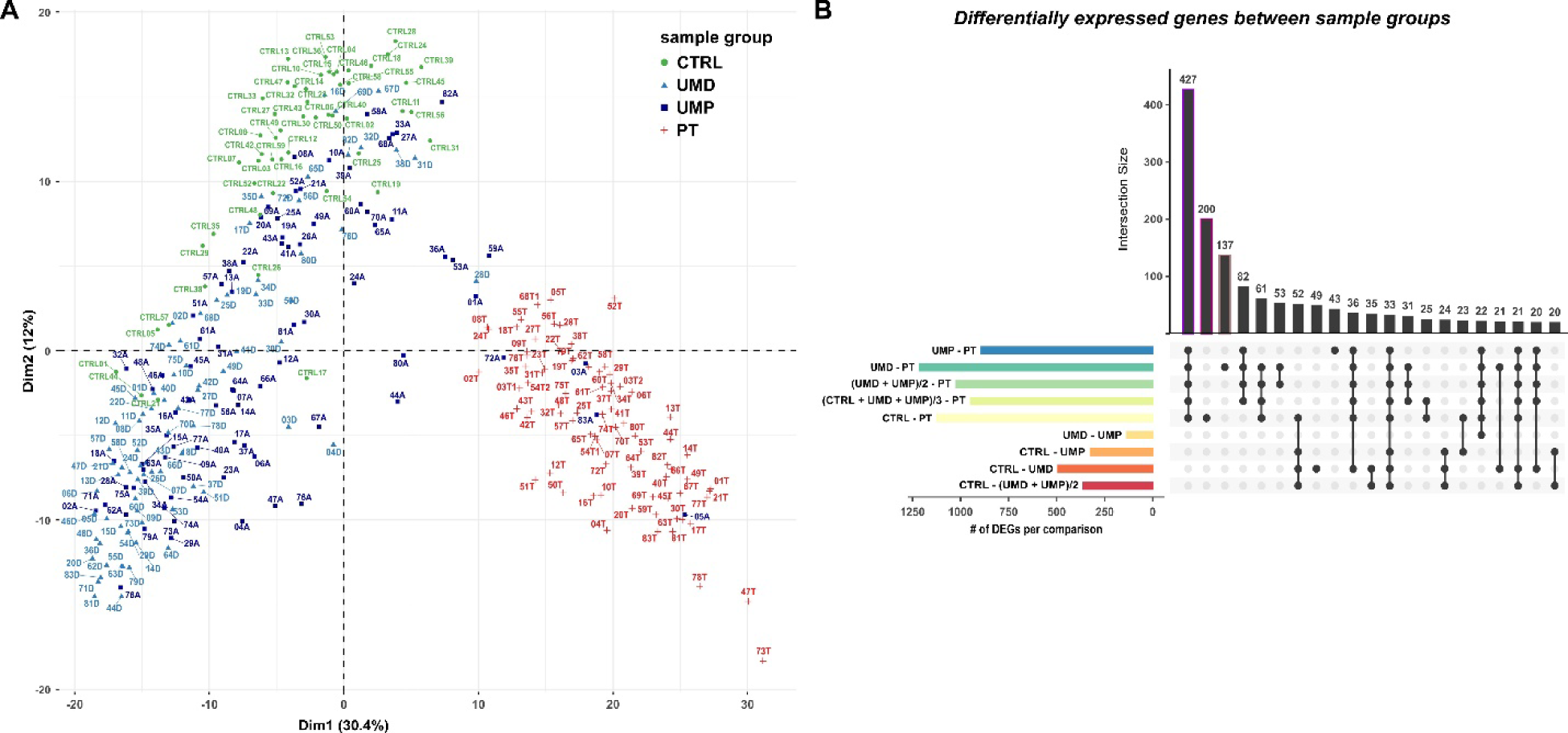
**A) Principal Component Analysis (PCA) of primary tumor (PT), uninvolved margin (UM) and control (CTRL) samples.** PCA was performed based on the expression of panel genes. UM samples include uninvolved margin proximal to primary tumor (UMP) and uninvolved margin distal from the primary tumor (UMD). This analysis illustrates the broad dispersion of breast cancer and morphologically normal tissue samples across the main principal axes. Each point represents the orientation of a sample projected into the transcriptional space, color-coded to indicate its group membership. Tumor profiles primarily aggregate in a distinct quadrant of the transcriptional space, whereas UM and CTRL tissues occupy two separate quadrants. The first PC represents the maximum variance direction in the data (30.4%), that corresponds to the differences between primary tumor versus all other samples. The second PC primarily reflects differences between UM and CTRL profiles, with proximal and distal profiles displaying a broad spread across the main principal axes. The PCA plot was generated with the fviz_pca_ind function from the R package factoextra (version 1.x0.7). **B) Differential gene expression analysis across primary tumor (PT), uninvolved margin (UM) and control (CTRL) samples.** UM samples include uninvolved margin proximal to primary tumor (UMP) and uninvolved margin distal from the primary tumor (UMD). The highest number of DEGs (false discovery rate (FDR) ≤0.05 and a log-fold change of ≥1; QLF test) is observed when comparing PT versus all non-malignant tissues, while UMP and UMD share few DEGs indicating overall similarity between their expression profiles. The variable contribution plots were generated with the fviz_contrib function.

Examination of the functional annotation of differentially expressed genes (DEGs) identified, as expected, enrichment of gene ontology terms and pathways previously associated with cancer (Figure S1). The observed sets of enriched biological terms and pathways among the primary tumor profiles, such as those related to proliferation and cell cycle, reflected the aggressiveness of those cancers and the unfavorable outcome of this cohort.

The second principal component of the PCA of all samples, showed heterogeneity within the non-malignant group (Figure 2A). CTRL samples formed a relatively homogeneous population, distinct from uninvolved margins (UMs), while UM samples were more variable and dispersed over a broader area. To investigate this further, we performed PCA solely on UM and CTRL samples. This revealed a subset of UMs forming a distinct population (Figure S2A). Notably, the list of the top 50 genes that accounted for the variability in the first principal component (y-axis), distinguishing between UM and CTRL samples, included genes related to epithelial matrix structure and organization (Figure S2B).

### UM tissues display abnormal features according to PAM50 gene classifier

AIMS and PAM50 predictors were applied to all samples to corroborate the histopathological classification. Both tools validated and classified all samples from reduction mammoplasty surgeries (CTRL) as normal-like. In tumor fragments, AIMS and PAM50 classifiers tend to agree more in assigning the basal-like subtype to PT samples, rather than Luminal A, Luminal B or HER2-enriched [AIMS vs. histopathological classification accuracy: 70% (Luminal A), 74% (Luminal B), 65% (HER2-enriched), and 88% (Basal-like) / PAM50 vs. histopathological classification accuracy: 0 (Luminal A), 66% (Luminal B), 70% (HER2-enriched), and 87% (Basal-like)] (Additional File 1, Supplementary Table 5). However, we noticed some disagreement in UM samples; while AIMS classified most as normal-like, PAM50 assigned ∼40% of the UM samples to tumor-like subtypes. This discrepancy could potentially be explained by the different number of genes incorporated and the different principles used by each tool. At the same time, PAM50 probability scores for individual samples indicated that classification of UMs often balanced between the normal-like and tumor-like subtypes suggesting the presence of features deviating from the “normal” state (Additional File 1, Supplementary Table 5).

### A distinct cluster emerges within uninvolved margin tissues

Through hierarchical clustering of all samples in our dataset, using the expression profiles of panel genes, we identified four distinct clusters (Figure 3). Two of these clusters, referred to as Cluster 1 and Cluster 2, were predominantly populated by PT samples. Clusters 1 and 2 appeared to be formed according to PT molecular subtypes, as determined by histopathological evaluations combined with AIMS and PAM50 predictors. Cluster 1 (n=25) predominantly contained HER2-enriched and basal-like tumors, whereas Cluster 2 (n=57) was mainly composed of luminal tumors. The remaining two clusters included non-malignant samples. CTRL samples, barring two exceptions, populated Cluster 3 (n=145), while UM samples dispersed between Clusters 3 and 4 (n=68). We further sought to investigate why UM samples split between these two clusters, while CTRL samples largely coalesced within Cluster 3, despite both originating from the same tissue type. The notable difference in age between breast cancer patients (UM samples) and individuals subjected to reduction mammoplasty surgeries (CTRL samples) could potentially be an element adding to the situation, although we included age as a covariate in Differential expression analysis.

**Figure 3.**
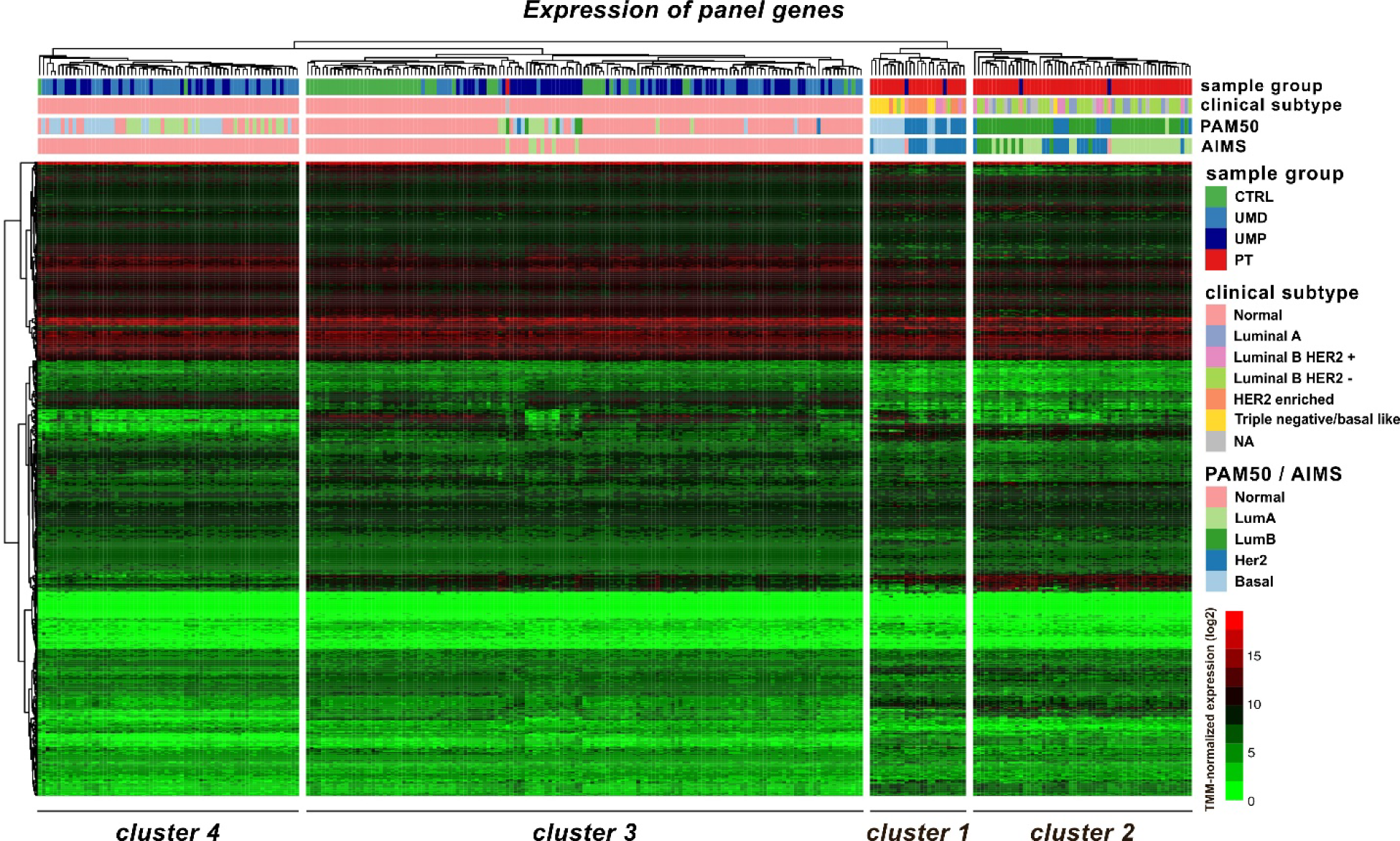
Hierarchical clustering of primary tumor (PT), uninvolved margin (UM) and control (CTRL) samples. Clustering performed using the expression profiles of panel genes reveals four distinct clusters. Cluster 1 (n=25) primarily contains HER2-enriched and basal-like tumors, whereas Cluster 2 (n=57) mainly includes luminal tumors. The remaining two clusters accommodate non-malignant samples. CTRL samples, with the exception of two cases, are mainly clustered in Cluster 3 (n=145), while UM samples are distributed across Clusters 3 and 4 (n=68). The clinical subtype assigned via histopathological examination is illustrated for all primary tumors. Additionally, subtype information, assigned via AIMS and PAM50 predictors is included for all samples in the dataset (Additional File 1, Supplementary Table 6).

A subsequent DE analysis that incorporated cluster assignment along with the sample type, revealed that the highest number of differentially expressed genes was observed between CTRL samples in Cluster 3 and UM samples in Cluster 4 (FDR ≤ 0.05, log-fold change of ≥1). The second-highest number of DEGs appeared when comparing UM profiles between Clusters 3 and 4 (Figure S3).

Remarkably, the top down-regulated genes in UM tissues in Cluster 4 included keratins: *KRT14, KRT15, KRT17, KRT6B, KRT5, KRT7, KRT19*, cell adhesion-related genes: *CDH1, CDH3, EPCAM*, and a matrix metallopeptidase *MMP7*. This list also comprised transcription factors *FOXI1, FOXA1* - tumor suppressor or candidate tumor suppressor genes, dual-role genes *RAB25, NRG1, SPDEF, TRIM29,* and the *GABRP* gene previously associated with breast cancer metastatic potential. A selection of these genes is presented in Figure 4A. The statistical significance of these findings persisted (p<0.05, Quasi-Likelihood F-test - QLF) when including the sample group information and even under multiple comparison scenarios (UMs in Cluster 4 versus UMs in Cluster 3, UMPs in Cluster 4 versus UMPs in Cluster 3, UMDs in Cluster 4 versus UMDs in Cluster 3) (Figure 4B). These genes exhibited a bimodal expression pattern in both types of UMs, best explained by the split of UM samples between Clusters 3 and 4. They form a distinct signature, hereby named as KAOS signature for Keratin-Adhesion-Oncogenes-Suppressors.

**Figure 4.**
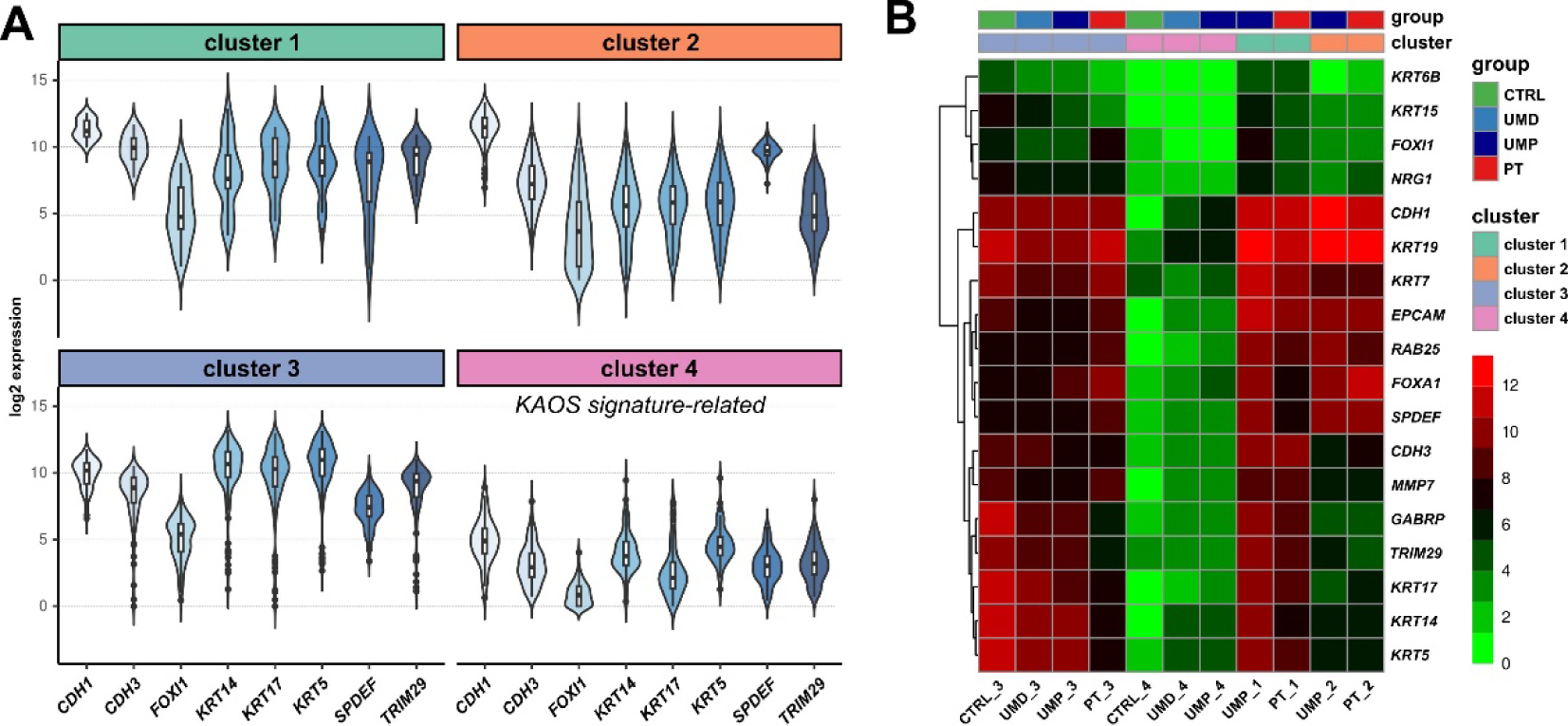
**A) Violin/box plots illustrating the expression profiles of genes included in the identified signature in identified Clusters 1, 2, 3, and 4.** Low expression of selected genes is noted in Cluster 4 compared to Cluster 3. **B) Heatmap with average expression of genes included in the identified signature in Clusters 1, 2, 3, and 4.** Further stratification, including sample group i.e. primary tumor (PT), uninvolved margin proximal (UMP) and distal (UMD) from the PT, and control (CTRL) highlights a relative down-regulation of gene expression in UMP, UMD, and CTRL samples located in Cluster 4 compared to those in Cluster 3.

### Cluster 4 is distinguished from other non-malignant profiles through Enrichment Analysis

Enrichment analyses were conducted to identify disrupted Gene Ontology terms and KEGG pathways in samples located in Cluster 4 (hypergeometric test, FDR < 0.05) (Additional File 1, Supplementary Tables 6, 7). Analyses revealed that primarily epithelial/stem cell developmental processes, the estrogen signaling pathway, and the cell adhesion molecules pathway were up-regulated in Cluster 3UMs relative to Cluster 4 UMs. In contrast, the "Regulation of Lipolysis in Adipocytes" and the "PPAR Signaling Pathway" were significantly down-regulated in Cluster 3 UMs compared to Cluster 4 UMs (Figure 5A). Notably, all above-described observations remained consistent when performing multiple comparisons between UM and CTRL samples located in Clusters 3 and 4 (Figure 5B).

**Figure 5.**
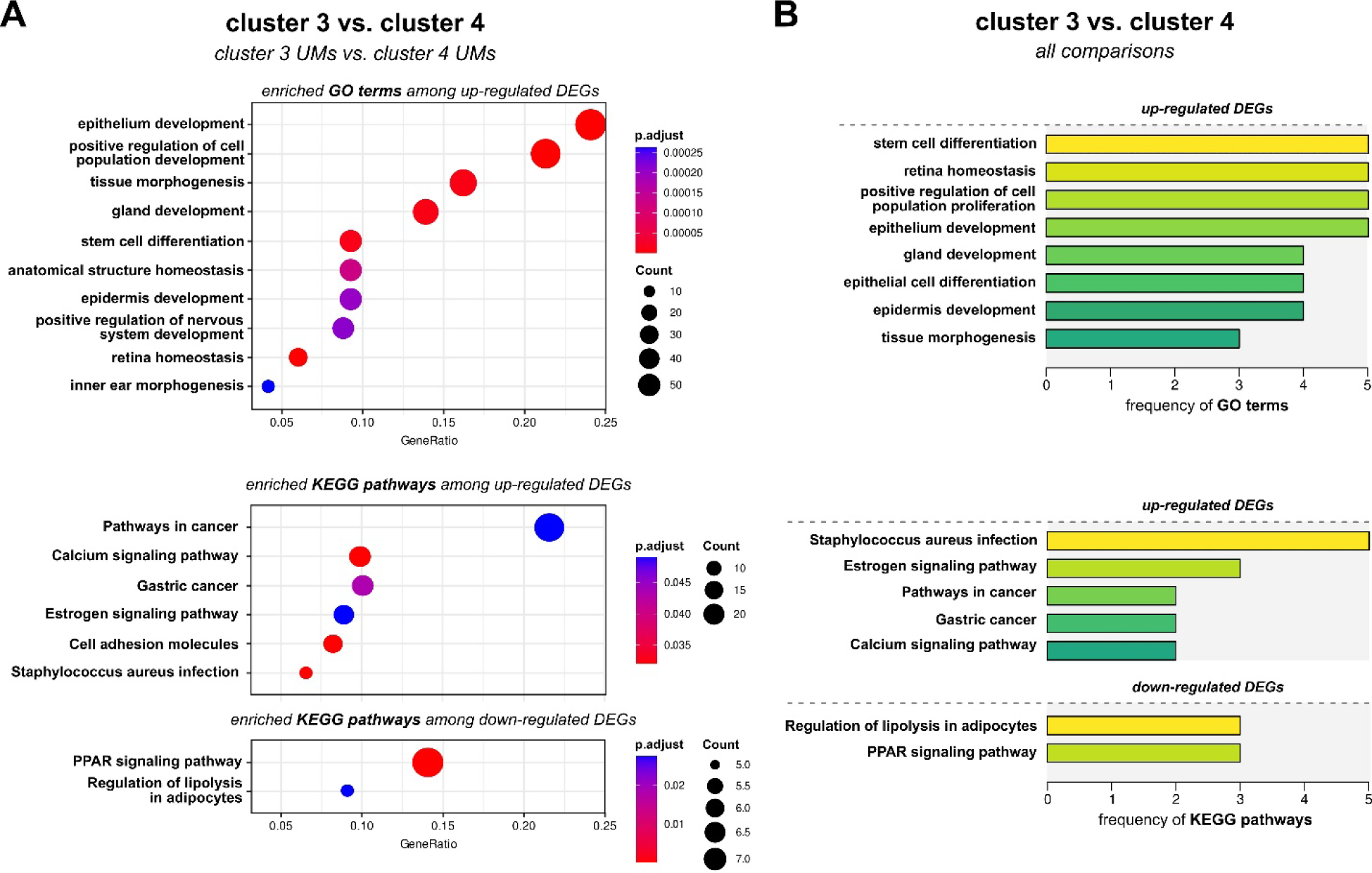
**A) Enrichment analysis across uninvolved margin (UM) samples in Clusters 3 and 4.** Analysis performed for differentially expressed genes (DEGs) identifies Gene Ontology (GO) terms and KEGG pathways. Maximum ten significantly enriched GO terms/KEGG pathways are presented here.The DEG sets were filtered to retain only those demonstrating a log fold change (logFC) of greater than or equal to 1, with an adjusted p-value (p adj) of less than 0.05. **B) Frequency of enriched features across multiple comparisons involving Clusters 3 and 4.** Comparisons include UM_3 vs. UM_4, UMP_3 vs. UMP_4, and UMD_3 vs. UMD_4, CTRL_3 vs. UMs_4; (full list of DE tests is available in the Additional File 1, Supplementary Tables 6,7). GO terms/KEGG pathways with frequency at least 3 among the conducted comparisons are presented. The Estrogen signaling pathway, the PPAR signaling pathway, and the Regulation of lipolysis in adipocytes pathway consistently appear enriched among DEGs upregulated in Cluster 4 UM samples upon multiple comparisons.

It should be noted here that the customized RNA-sequencing panel’s ability to efficiently capture crucial information, representative of the full mammary tissue transcriptome, was validated by analyzing two distinct external datasets (full transcriptome – custom RNA-seq panel) of PT and paired UM samples from the same 18 breast cancer patients that originated from other breast cancer study in our lab (Figure S4).

### Cluster 4 significantly associates with patients’ clinical outcome

Interestingly, Cluster 4 shared a greater degree of similarity with Clusters 1 and 2, which were dominated by malignant profiles, compared to cluster 3, populated by CTRLs and UMs (Figure S5). Cluster 4 was significantly enriched with UM samples (p=1.98e-08, Fisher’s test), encompassing 23% of all samples and 40% of total UM samples, representing 41% of patients. Furthermore, it was significantly enriched (p=7.62e-05, Fisher’s test) with samples that were classified by PAM50 as one of the breast cancer molecular subtypes (when comparing within UMs only). This association was significant for all UMs, as well as for both distal and proximal UMs (p= 1.02e-08 and p= 0.00256, respectively).

Identification of a distinct patient group, having both UM samples (proximal and distal) in one cluster, allowed us to execute comprehensive comparisons using follow-up information collected for each patient. Patients with both UMs in Cluster 4, as opposed to Cluster 3, exhibited smaller tumor sizes as measured by ultrasonography and pathological examination (p=0.0013 and p=0.033, respectively, Mann-Whitney U test) and were older (p=0.025, Mann-Whitney U test). Cluster 4 membership was also associated with tumor HER2 positive status (p= 0.004265, Fisher’s test). Upon restricting our analysis to patients with only one UM sample assigned to either Cluster 3 or 4, it was observed that Cluster 4 had a notable overrepresentation of patients with a positive death status (p=0.04493345 and p=0.01512627, Fisher’s test for UMD and UMP, respectively). Finally, when performing another comparison, for patients with a strictly defined UMs clustering pattern, i.e., UMD in Cluster 3 and UMP in Cluster 4, a substantial link to patient death status emerged, contrasting with the patients who had the reverse UM assignment (UMD in Cluster 4 and UMP in Cluster 3) (p= 0.001396). These findings indicate that the spatial information of uninvolved mammary tissue, obtained by samples at different distance from the primary tumor, could reveal different pieces of information of the patient’s clinical picture.

## DISCUSSION

This study takes an important step towards understanding the properties of microscopically normal, uninvolved mammary tissue in breast cancer patients with unfavorable outcomes. Our results provide new insight into the concept of biological abnormality in histologically normal mammary gland tissue, removed at different distances from the primary tumor ^11–13,27^.

We identified a distinct subset of histologically non-cancerous uninvolved mammary (UM) samples, referred to as Cluster 4, which demonstrates unique attributes in contrast to other non-cancerous UM samples and, importantly, to mammary gland samples collected from individuals without cancer (CTRL). Notably, Cluster 4 is significantly enriched with UMs (both proximal and distal) categorized as tumor-like by PAM50. This likely suggests the presence of feature characteristics in Cluster 4 UMs divergent from the “normal” state.

Furthermore, a distinct gene signature present in Cluster 4 UMs, named as KAOS signature, involving the down-regulation of genes participating in various processes, supports the above-mentioned thesis. Genes comprising this signature can be grouped into two main categories; a) cell adhesion and structural support genes, and b) transcription factors and tumor suppressors/oncogenes. The first group included genes encoding for keratins (*KRT14, KRT15, KRT17, KRT6B, KRT5, KRT7, KRT19*), metallopeptidases (*MMP7*), and cell adhesion molecules (*CDH1, CDH3*, *EPCAM*). Motility keratins *KRT5, KRT14*, and *KRT17,* have been previously implicated in “subtype switching”, i.e. switching of molecular subtype between lung and pleura metastases versus the primary breast tumor ^28^, as well as deemed essential for the tumorigenic potential and migration of a basal-like breast cancer cell line along with *GABRP* ^29^. Furthermore, *KRT19, KRT7*, and *KRT15* are individually linked to breast cancer, especially associating with metastatic potential and poor patient outcome ^30–32^. Diminished cytokeratin, cell adhesion-related (*CHD1, CDH3, EPCAM*) and matrix metallopeptidase (*MMP7*) gene expression indicates the presence of alterations linked to invasiveness and signifies disruptions in the cytoskeleton, pointing to loss of adhesion and epithelial tissue integrity. Subsequently, these observations point to epithelial-to-mesenchymal transition (EMT), an otherwise normal process during which epithelial cells acquire migratory and invasive properties, observed also in tumor metastasis ^33,34^. Nonetheless, we did not observe concomitant up-regulation of mesenchymal markers. This suggests the potential presence of a partial/hybrid EMT ^35^. The second group contained candidate oncogenes and candidate tumor suppressor genes in breast cancer (*RAB25, NRG1, SPDEF*), reportedly having dual-functioning roles associated with estrogen (ER) status ^36–38^. Additionally, this group included the tumor suppressor *TRIM29* gene, whose depletion has been linked with preneoplastic changes such as loss of polarity and increased migration and invasion in non-tumorigenic breast cells ^39^, as well as alteration of keratin expression to enhance cell invasion in squamous cell carcinoma ^40^. Lastly, members of the forkhead box transcription family (*FOXA1, FOXI1)* previously associated with breast cancer ^41,42^, were part of this group. The observed down-regulation of genes with dual roles in cancer, both established and candidate tumor suppressor genes, as well as transcription factors, underscores the disturbed environment in the Cluster 4 UM samples.

Interestingly, Cluster 4 was further characterized by the down-regulation of the estrogen signaling pathway and the up-regulation of the “Regulation of lipolysis in adipocytes’’ and the “PPAR signaling” pathways. Deregulation of the latter, an upstream effector of fatty acid oxidation ^43^, suggests a link to metabolic imbalance. Disruption of metabolic-related processes has been observed in patients with altered PPAR signaling, reinforcing the established role of PPARs in lipid transport, fatty acid oxidation, and their involvement in crosstalk with other lipogenic pathways ^44^. Intriguingly, PPARs can share common ligands with estrogen receptors (ERs) and both have contrasting regulatory effects on the PIK3K/AKT signaling pathway, which influences breast cancer cell survival and proliferation ^45^.

We observed a significant association between cluster membership and less favorable patient outcomes, as patients with UM samples in Cluster 4 were associated with a positive death status, and also with smaller size tumors. The latter finding is intriguing, although not straightforward to explain. Patients enrolled in this study, barring two exceptions, experienced mortality primary attributed to breast cancer itself, disease recurrence, or the emergence of secondary tumors. However, the presence of comorbidities (undocumented here) such as hypertension, cardiovascular disease (CVD), and type 2 diabetes can affect the progression of the disease, complicate treatment and influence the patient’s health outcome ^46^. Although we did not find a direct significant association between cluster membership (for UM samples) and recurrence or secondary tumor events in corresponding patents, our findings could likely reflect the overall systemic aggressiveness of the disease. The concept of untransformed cells dissociating from the original diseased organ, disseminating via the vascular system, and incorporating into parts of otherwise normal-appearing organs to seed metastases, has been recently raised again through the work of Rahrmann et al ^47^. The disturbed tissue microenvironment found in Cluster 4 UMs could potentially facilitate homing of these untransformed cells early on, thus assisting in the spread of cancer throughout the body and ultimately leading to death.

The etiologic field theory ^48^, a different perspective on the field effect theory initially proposed by Slaughter’s group in 1953 ^49^, supports the above-mentioned thesis. This concept embraces tumor-host and gene-environment interactions and highlights the existence of an abnormal tissue microenvironment present within microscopically normal tissue that can influence every stage of tumor development. Importantly, the etiologic field effect concept challenges the notion that markers exclusively indicate neoplasia. Instead, it suggests that these markers may represent environmental changes, including the potential contribution of non-transformed cells and extracellular matrices to neoplastic evolution. A continuous model, involving multiple stages, favoring the acquisition of alterations, might be a better representation of a realistic tumorigenic process.

Our findings present an alternative perspective to a previous study, which postulated that histologically normal tissue adjacent to breast cancer exhibits only minimal gene expression changes compared to breast reduction tissue ^50^. According to that study, these differences in gene expression primarily represented individual tissue- and patient-specific variability, rather than any associations with the patient’s clinical picture. However, it is worth noting that our study differed in terms of patient selection, as we focused on patients with adverse outcomes, including the presence of recurrence, the emergence of a second independent tumor, or mortality, as the principal inclusion criteria. Furthermore, the determination of tumor adjacency varied across these two studies. Consequently, our findings indicate the development of a pre-tumoral, change-favoring environment, a feature characteristic of patients with a higher risk of recurrence and a decreased survival rate.

While our study offers valuable insights into the molecular changes occurring within the uninvolved mammary gland of breast cancer patients with unfavorable outcomes, it does come with certain limitations. We understand that we may not have captured the complete spectrum of molecular changes happening within these tissues, since we only focused on aberrations at the gene expression level. Secondly, we did not include samples from metastatic sites in our study design. The incorporation of samples from secondary lesions would have allowed for a thorough assessment and comparison of gene expression profiles among primary tumor sites, surrounding normal-appearing gland tissue, and metastatic sites. In this context, examining the tumor microenvironment, specifically focusing on alterations in stromal and immune cells, might have provided valuable insights. However, the procurement of the corresponding samples presents significant challenges. Finally, the transition from a cross-sectional to a longitudinal study design might have allowed us to better track the evolution of the tissues over time, their contribution to cancer progression and metastasis as well as the influence of external factors, such as the patient’s lifestyle and environmental factors.

Nevertheless, the significant link between the clustering pattern and patient death status implies a potential prognostic value, suggesting that the spatial distribution of uninvolved mammary tissue could hold crucial information about breast cancer outcomes.

Our study highlights the potential presence of a pre-tumorigenic environment, within the ostensibly normal mammary gland tissue, promoting changes that are closely linked to patient mortality. The aberrant gene expression profiles of uninvolved mammary tissue intriguingly exhibit tumor-like characteristics as shown by the PAM50 predictor, marked by dysregulation of crucial pathways such as estrogen and PPAR signaling.

It remains to be determined whether these observed alterations stem from the nearby tumor’s influence or signify the independent emergence of early pre-tumorous conditions facilitated by a perturbed environment. The strong association of Cluster 4 characteristics with mortality, but not directly with recurrence, may suggest these features are more indicative of the disease’s systemic aggressiveness than of its potential to re-emerge.

This study offers an indication for comprehensive monitoring of breast cancer patients with recurrence or secondary tumor events. Integrating molecular assessments of non-malignant mammary tissue into disease management strategies could enhance personalized patient care, including improved survival prediction.

## Abbreviations

BCS: Breast-conserving surgery
PT: Primary tumor
UMP: Uninvolved margin proximal to primary tumor
UMD: Uninvolved margin distal from primary tumor
CTRL: control
DE: Differential expression
DEGs: Differentially expressed genes
EMT: epithelial-to-mesenchymal transition
PCA: Principal component analysis

## Authors Contributions

Conceptualization: A.P., J.P.D., N.F., M.A. Resources: J.P.D., T.N., W.Z., Ł.S., M.J., J.J., E.Ś., M. L-J., D.B., J.H.,M.N., R.P., J.S., J.H., H.D., B.B-O. Data Curation: N.F., A.P., M.A., T.N., K.D. Investigation: M.A., N.F., K.D., U.Ł., K.C., M.H. Methodology: A.K., N.F., M.H. Data analysis: M.J., J.M. Visualization: M.J, M.A. Interpretation: M.A., A.P., M.J., J.M., A.K. Manuscript writing-original: M.A., M.J., A.P. Manuscript writing – review and editing: A.M., M.J., A. P., J.P.D., J.M., N.F., A.K., M.H., H.D. All authors have read and approved the manuscript. M.A. and M.J. have contributed equally to this work. Supervision: J.P.D., J.M., A.P. The work reported in the paper has been performed by the authors, unless clearly specified in the text.

## Ethics approval and consent to participate

Tissue samples and patient histories were provided for this study by the Oncology Centre in Bydgoszcz and the University Clinical Centre in Gdańsk, who, under a research protocol approved by the Bioethical Committee at the Collegium Medicum, Nicolaus Copernicus University in Toruń (approval number KB509/2010) and by the Independent Bioethics Committee for Research at the Medical University of Gdansk (approval number NKBBN/564/2018 with multiple amendments), recruited and enrolled all donors under informed and written consent, collected, and stored all tissue samples.

## Consent for publication

Not applicable.

## Data availability

The bulk RNA-seq data generated in this study are available from the European Genome-Phenome Archive (EGA, https://ega-archive.org/) under ID EGAS50000000011 accession number. All other primary data analyzed and presented in this study are located in the Supplementary files attached to this manuscript.

## Corresponding author

Correspondence and requests for materials should be addressed to Arkadiusz Piotrowski.

## Acknowledgements

The authors wish to thank Agata Wojdak and Kinga Drężek for their help with administrative and wet lab activities, respectively. We also would like to thank all the patients and volunteer controls for acceptance to participate in the study and sample contribution; hospital staff involved in the patient recruitment process in Oncology Center - Prof. Franciszek Łukaszczyk Memorial Hospital in Bydgoszcz and University Clinical Centre in Gdańsk. Lastly, we thank Dr. Leszek Kalinowski for the access to selected laboratory facilities.

## ADDITIONAL FILES

**Additional File 1.xls: Supplementary Tables 1,2,3,4,5,6,7**

**Supplementary Table 1. Clinicopathological characteristics of breast cancer patient cohort**. Primary tumor (PT) samples were collected from 83 individuals diagnosed with breast cancer. Cancer TNM stage (a) and tumor grade (b) information. (c) Histological types of breast tumors recognized: invasive ductal carcinoma (IDC), invasive lobular carcinoma (ILC), neuroendocrine, mucinous and papillary carcinoma. Comedo refers to comedocarcinoma, a type of pre-invasive breast neoplasia. (d) In situ: ductal carcinoma in situ (DCIS), lobular carcinoma in situ (LCIS). (e) BCT-breast-conserving therapy; SLND - sentinel lymph node dissection; MRM (Patey)- Modified Radical Mastectomy (Patey); Q - breast quadrantectomy (included in Breast-conserving therapy procedures); ALND - axillary lymph node dissection; LS - lymphoscintigraphy; M - mastectomy. (f) Estrogen receptor (ER), (g) progesterone receptor (PR), (h) Ki67 proliferation marker, and (i) HER2 status were assessed by Immunohistochemistry (ICH). (j) ER, PR, HER2, and Ki67 scores were used to assign tumors to biological subtypes. (k) tumor size measured by Ultrasonography. (l) Collected samples: PT1 - primary tumor 1, PT1 - primary tumor 2 (PT1&PT2 refer to two different samples from two different areas of multifocal primary tumor), UMP - uninvolved margin proximal to the primary tumor (> 1 cm and always in shorter distance than corresponding UMD), UMD - uninvolved margin distal from the primary tumor (1,5 - 5 cm).

**Supplementary Table 2. Age of individuals subjected to reduction mammoplasty surgeries, serving as controls**. CTRL – control.

**Supplementary Table 3. Target genes included in the in-house designed panel and information regarding internal groups.** (a) Gene symbols of targets included in panel, (b) Ensembl ID, (c) Ensembl ID as in Gencode (version 35), (d) Gene symbol according to the HUGO Gene Nomenclature Committee, (e) EntrezID, (f) Functional classification.

**Supplementary Table 4. Sequencing coverage and statistics of each sample included in our dataset.**

**Supplementary Table 5. Hierarchical clustering of samples and assignment of subtypes by AIMS and PAM50.** (a) Sample category: T - Tumor, D - Distal, P - Proximal, CTRL – Control. (b) Combined Individual ID and sample category. (c) Cluster classification produced by Hierarchical clustering. (d) Combined sample category and cluster information. (e) Subtype assigned by histopathological evaluation – all uninvolved margin and control samples were classified as “Normal”. (f) AIMS and (g) PAM50 predictors assigned subtypes to all samples included in the dataset: Her2 – Her2 enriched, LumA – Luminal A, LumB – Luminal B, Basal – Basal-like. (h) Probability scores of assigned subtypes by AIMS and PAM50.

**Supplementary Table 6. Gene Ontology (GO) terms identified via enrichment analyses, including cluster membership information.** Enrichment analysis was conducted for DEGs identified by comparing primary tumor (PT), uninvolved margin proximal to primary tumor (UMP), uninvolved margin distal from the primary tumor (UMD), and control (CTRL) samples. (a) Unique identification number and (b) short description of identified Gene ontology (GO) terms. (c) Ratio of genes included in the panel, associated with the particular GO term. (d) Ratio of background genes (full transcriptome) associated with the particular GO term. Enrichment (e) p value, (f) adjusted p value and (g) q value. (h) Genes associated with the particular GO term. (i) Direction of enrichment for the particular GO term; up - upregulation, down - downregulation. (j) Information regarding the comparison between different groups; PT - primary tumor, UMP - uninvolved margin proximal to the primary tumor, UMD - uninvolved margin distal from the primary tumor, CTRL - control. Numbers 1,2,3,4 indicate cluster membership (Figure 3).

**Supplementary Table 7. KEGG pathways identified via enrichment analyses, including cluster membership information.** Enrichment analysis was performed for DEGs identified by comparing primary tumor (PT), uninvolved margin proximal to primary tumor (UMP), uninvolved margin distal from the primary tumor (UMD), and control (CTRL) samples. (a) Unique identification number and (b) short description of identified KEGG pathways. (c) Ratio of genes included in the panel, associated with the particular KEGG pathway. (d) Ratio of background genes (full transcriptome) associated with the particular KEGG pathway. Enrichment (e) p value and (f) q value. (g) Direction of enrichment for the particular KEGG pathway; up - upregulation, down - downregulation. (h) Information regarding the comparison between different groups; PT - primary tumor, UMP - uninvolved margin proximal to the primary tumor, UMD - uninvolved margin distal from the primary tumor, CTRL - control. Numbers 1,2,3,4 indicate cluster membership (Figure 3).

## SUPPLEMENTARY FIGURES

**Figure S1. A) Gene Ontology (GO) terms and B) KEGG pathways identified across primary tumor (PT), uninvolved margin (UM), and control (CTRL) samples.** Enrichment analysis performed for Differentially expressed genes (DEGs)identified GO terms and KEGG pathways enriched by comparing PT, uninvolved margin proximal to primary tumor (UMP), uninvolved margin distal from the primary tumor (UMD), and CTRL samples. The DEG sets from the QLF test were filtered to retain only those demonstrating a log fold change (logFC) of greater than or equal to 1, with an adjusted p-value (p adj) of less than 0.05. When applicable, CTRL samples were set as the baseline for Differential expression (DE) analyses, thus down-regulation corresponds to lower expression in CTRL vs. PT and alike. Mitotic division-related GO terms and the Cell cycle KEGG pathway, are amongst the enriched terms for down-regulated genes in PT samples versus non-malignant samples (UM + CTRL).

**Figure S2. A) Principal Component Analysis (PCA) of uninvolved margins and controls based on the expression of panel genes**. This analysis illustrates the broad dispersion of morphologically normal tissue samples (UMP, UMD) and samples from control individuals (CTRL) across the main principal axes. Each point represents the orientation of a sample projected into the transcriptional space, color-coded to indicate its group membership. The non-malignant tissues, encompassing uninvolved margin (UM) and control (CTRL) samples, display substantial heterogeneity within their group. The PCA plot was generated with the fviz_pca_ind function from the R package factoextra (version 1.x0.7) **B) Variability in the 1st principal component (expressed in percentages) within the nonmalignant samples.** Top 50 genes are included. The red dashed line in the graph indicates the expected average contribution. If the contribution of the variables were uniform, the expected value would be 1/length(variables). For a given component, a variable with a contribution larger than this cutoff could be considered as important in contributing to the component. Function get_pca_var, from the R package factoextra (version 1.x0.7) was used to extract the information on the contribution of genes to the observed variance and their correlation with the given principal component (PCA axis)

**Figure S3. A) Differential Expression (DE) analysis across nonmalignant samples.** The greatest number of differentially expressed genes (DEGs) is observed when comparing control (CTRL) samples in Cluster 3 to uninvolved margin (UM) samples in Cluster 4. Similarly, other comparisons corresponding to cluster 3 vs. cluster 4 contribute to most of the identified unique DEGs. Significantly deregulated genes were filtered to retain only those demonstrating a log fold change (logFC) of greater than or equal to 1, with an adjusted p-value (p adj) of less than 0.05.

**Figure S4. Analysis of two external datasets (full transcriptome, panel), including primary tumor (PT) and paired uninvolved margin (UM) samples.** Both datasets consisted of samples derived from the same 18 patients. **Principal Component Analysis (PCA) of PT and UM samples based on A) the expression of full transcriptome, B) subset of the full transcriptome dataset using the custom panel genes only, and C) the expression of the second dataset with custom RNA-seq panel, exhibit similar results.** These analyses illustrate the dispersion of breast cancer and morphologically normal tissue samples across the main principal axes. Each point represents the orientation of a sample projected into the transcriptional space, color-coded to indicate its sample group membership. Tumor profiles and uninvolved margin samples aggregate in distinct areas of the transcriptional space in all cases. **D) Differential Expression analysis across primary tumor (PT) and uninvolved margin (UM) samples based on the expression of full transcriptome or only the expression of genes from the custom RNA-seq panel.** The highest number of differentially expressed genes (DEGs) is revealed being down-regulated in uninvolved margin compared to primary tumor samples. The direction of expression change remains stable in both comparisons (transcriptome, panel). DEGs were filtered to retain only those demonstrating a log fold change (logFC) of greater than or equal to 1, with an adjusted p-value (p adj) of less than 0.05. **E) Enrichment analysis for DEGs identified when comparing uninvolved margin (UM) vs. primary tumor (PT) samples.** Analysis identifies shared enriched KEGG pathways despite using only a subset of genes in the custom RNA-seq panel.

**Figure S5. Heatmap of correlation-based distances between individual samples, sample groups and clusters.** Clusters pictured were identified through hierarchical clustering of panel genes expression. Colors correspond to distances represented as (1 - Pearson correlation) metric, with dark blue showing lowest sample distance (similar), and dark red the highest distance (dissimilar). Cluster 4 shows the highest distance with Clusters 1 and 2, which predominantly contain PT samples, as compared to Cluster 3, encompassing primarily UM and CTRL samples.

