## Supplementary figures and images for "Prelude to Malignancy: A Gene Expression Signature in Normal Mammary Gland from Breast Cancer Patients Suggests Pre-tumorous Alterations and Is Associated with Adverse Outcomes"

### Figure_S1

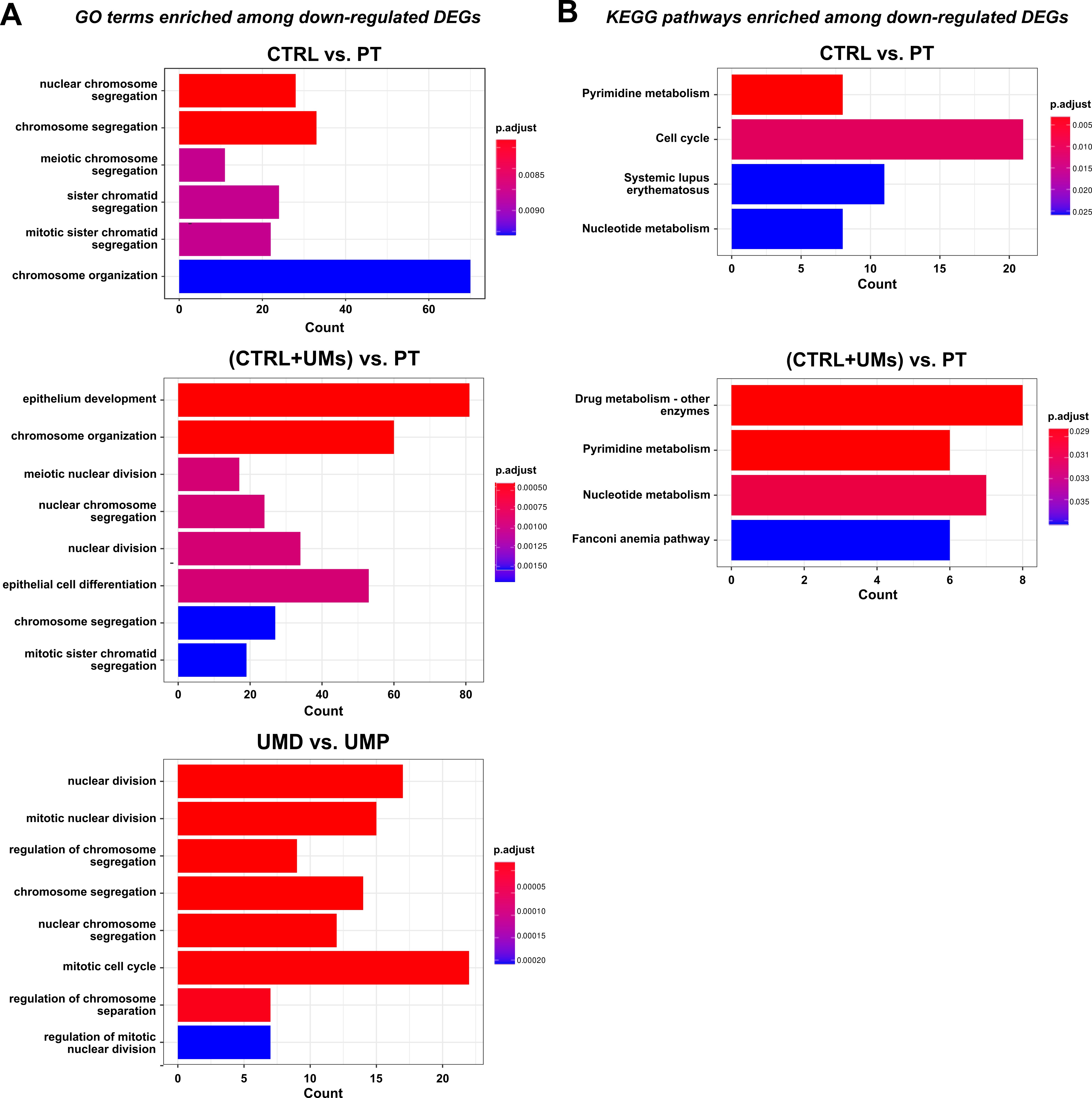

### Figure_S2

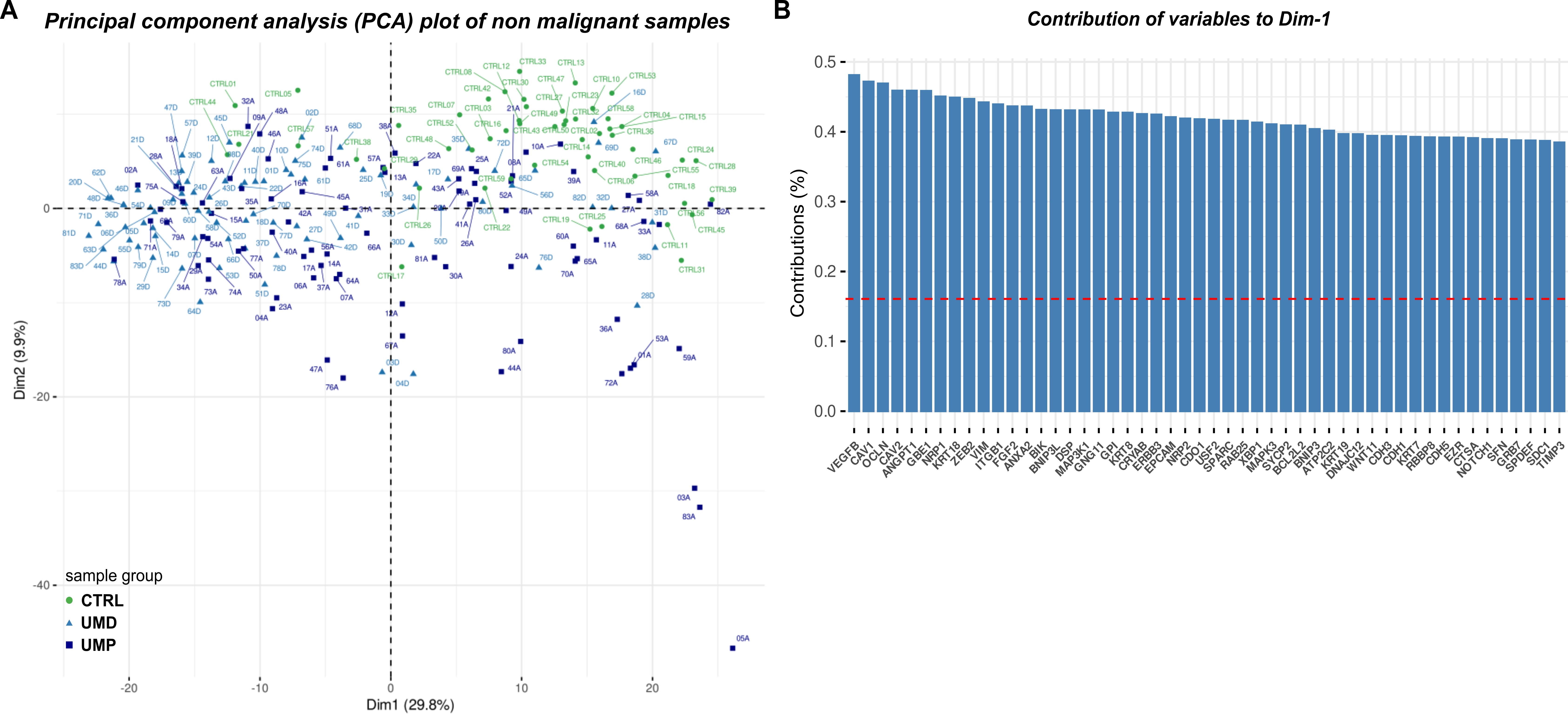

### Figure_S3

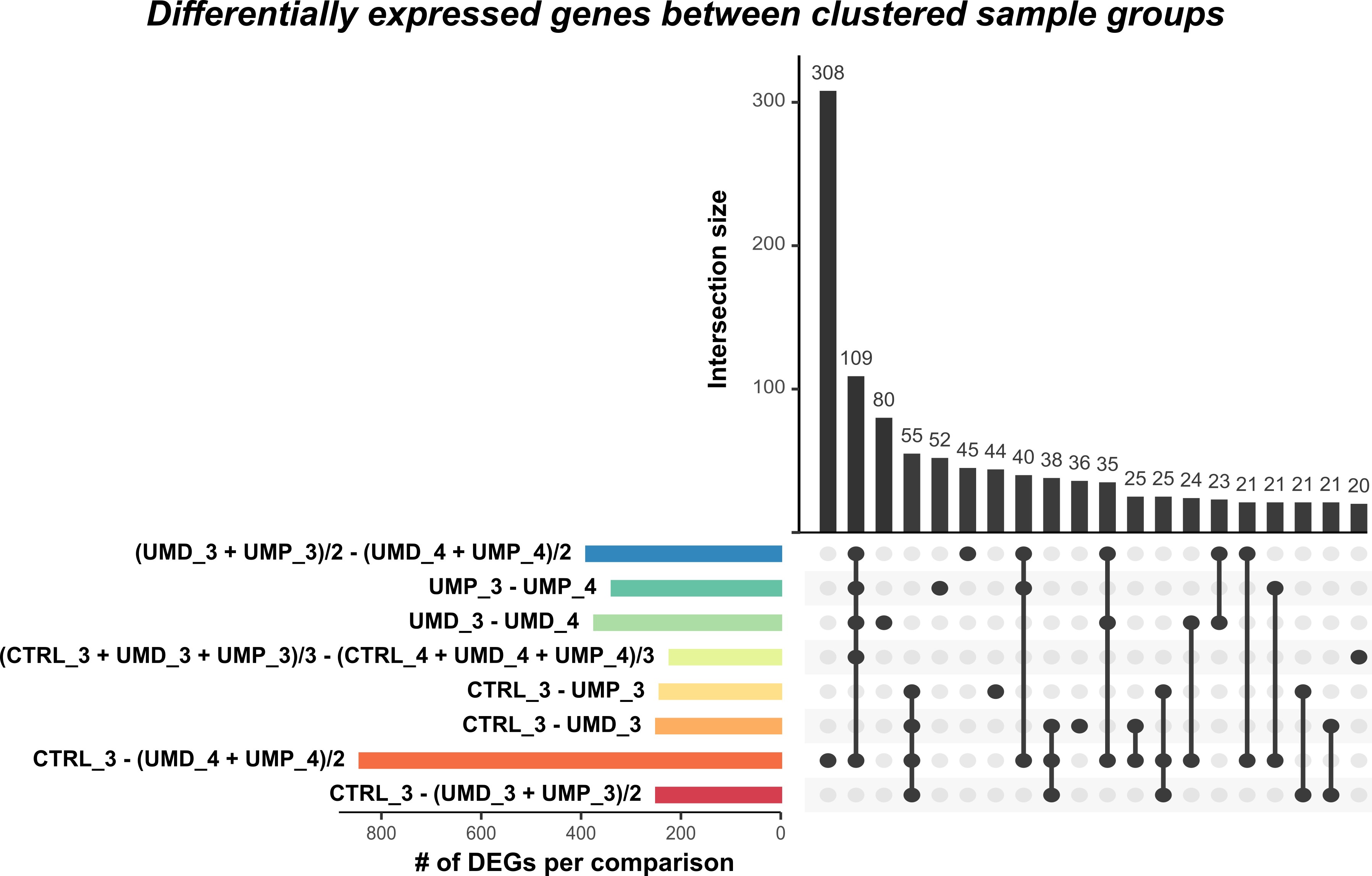

### Figure_S4

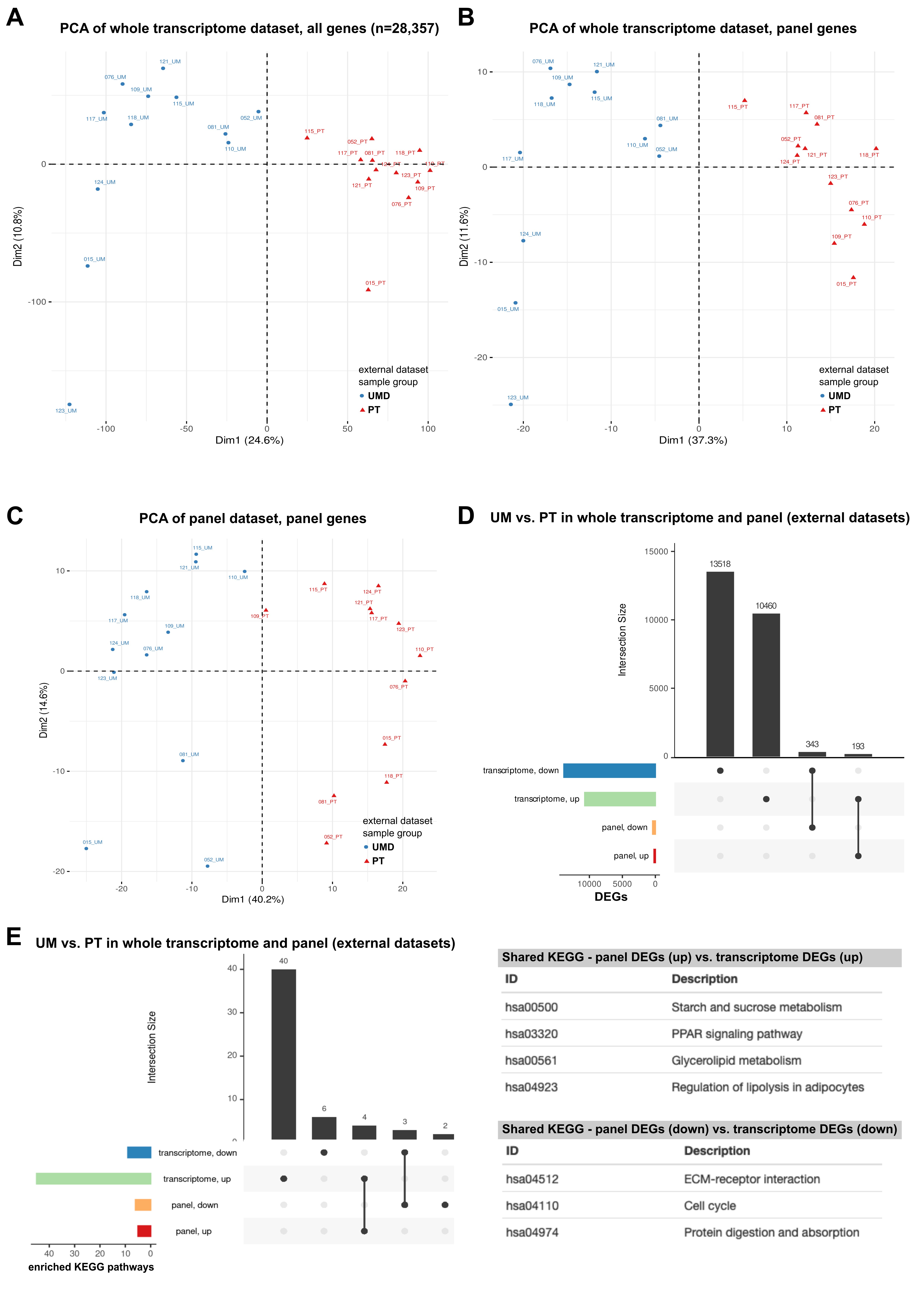

### Figure_S5

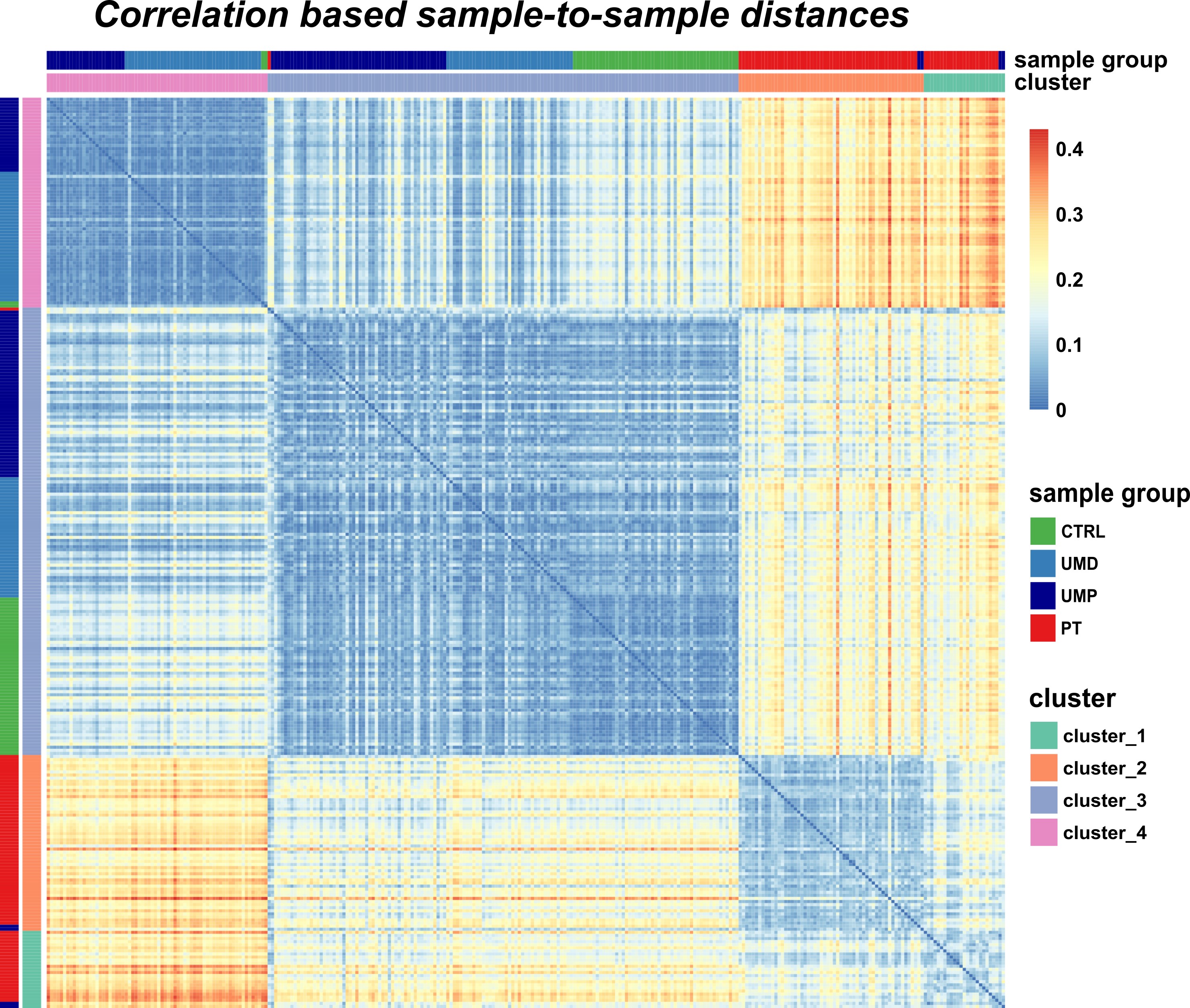
